# The structural basis of huntingtin (Htt) fibril polymorphism, revealed by cryo-EM of exon 1 Htt fibrils

**DOI:** 10.1101/2021.09.23.461534

**Authors:** Sergey Nazarov, Anass Chiki, Driss Boudeffa, Hilal A. Lashuel

**Affiliations:** Laboratory of Molecular and Chemical Biology of Neurodegeneration, School of Life Sciences, Brain Mind Institute, Ecole Polytechnique Fédérale de Lausanne (EPFL), CH-1015 Lausanne, Switzerland; BioEM Core Facility and Technology Platform, Ecole Polytechnique Fédérale de Lausanne (EPFL), CH-1015 Lausanne, Switzerland

**Keywords:** Huntington’s disease, Huntingtin, Fibrils, Cryo-EM, Polymorphism

## Abstract

The lack of detailed insight into the structure of aggregates formed by the huntingtin protein has hampered efforts to develop therapeutics and diagnostics targeting pathology formation in the brain of patients with Huntington’s disease. To address this knowledge gap, we investigated the structural properties of in vitro generated fibrils from exon1 of the huntingtin protein by electron cryo-microscopy and single-particle analysis. We show that wildtype and mutant exon1 of the huntingtin protein form non-helical fibrils with a polygultamine amyloid core composed of β-hairpins with unique characteristics that have not been previously observed with other amyloid filaments. The stacks of β-hairpins form long planar β- sheets (protofilaments) with variable stacking angle and occasional out-of-register state of individual β-hairpins. These features and the propensity of protofilament to undergo lateral association results in a high degree of fibril polymorphism, including fibrils composed of varying numbers of protofilaments. Our results also represent the first direct observation of how the flanking domains are organized around the polyglutamine core of the fibril and provide insight into how they might affect huntingtin fibril structure, polymorphism, and stacking of β-hairpins within its core structure. Removal of the first 17 amino acids at the N-terminus resulted in surprising intra-fibril structural heterogeneity and reduced fibril’s propensity to lateral associations. Overall, this work provides valuable insights that could guide future mechanistic studies to elucidate the sequence and structural determinants of huntingtin aggregation, as well as cryo-EM and structural studies of fibrils derived from huntingtin proteins and other disease-associated polyglutamine-containing proteins.

## Introduction

Huntington’s disease (HD) is the best-known disorder of a family of CAG repeat expansion neurodegenerative disorders. HD occurs when the polyglutamine (polyQ) tract in the first exon (exon 1) of the huntingtin protein (Htt) expands above a length of 36Q. Although the exact mechanisms underpinning the pathogenesis of HD are not yet completely understood, converging evidence points to Htt aggregation and accumulation in the form of nuclear and occasionally cytoplasmic inclusions as the central processes responsible for neurodegeneration in HD. Consistent with this hypothesis, Htt proteins with > 36 polyQ repeats are highly prone to misfold, aggregate, and form insoluble β-sheet-rich fibrillar aggregates^1–5^. Furthermore, the extent of aggregation in vitro and the degree of inclusion formation in cells are directly proportional to the length of the polyQ tract and are inversely correlated with disease onset age and severity, although there are some exceptions^6–9^. In animal models of HD, overexpression of N-terminal fragments containing expanded polyQ repeats induced inclusion formation, neurodegeneration, and HD-like phenotypes^10–15^. Furthermore, mutant Htt aggregation has been shown to result in the sequestration of wild-type Htt and other functional proteins, including chaperones and transcription factors, suggesting that Htt aggregation may contribute to the pathogenesis of HD through a combination of gain- and loss-of-function mechanisms^9,16,17^. Therefore, inhibiting mutant Htt aggregation or promoting the disassembly or clearance of Htt aggregates would represent a viable therapeutic strategy to treat HD and enable the testing of the aggregation hypothesis in HD. However, the lack of detailed insight into the structure of Htt aggregates has hampered efforts to develop effective Htt aggregation inhibitors or Htt-aggregate-specific binders that could be further developed into PET tracers for monitoring and quantifying pathology formation and development in HD brains.

Recent advances in electron cryo-microscopy (cryo-EM) have paved the way for solving the filament structures of several amyloid proteins prepared in cell-free systems or isolated from postmortem human tissues^18–22^. These structures have provided unprecedented insight into the structural basis of amyloid fibril formation and polymorphism associated with several systemic and neurodegenerative diseases. However, the structure of fibrils derived from proteins or peptides containing expanded polyQ repeats, e.g., polyQ peptides, Htt fragments or polyQ-repeat-containing amyloidogenic proteins, remains elusive.

Wild-type Htt is a large multidomain protein with functions in various cellular processes, and its structure was recently solved by cryo-EM^23^. Almost a quarter of the sequence of Htt, including exon 1, which comprises the polyQ segment mutated in HD and loops connecting structured domains, remains invisible due to high local flexibility. Current models of Htt aggregation and polyQ-dependent filament formation are based on X-ray fiber diffraction, electron paramagnetic resonance (EPR), solid-state NMR (ssNMR) and molecular dynamics studies^24–36^. X-ray powder diffraction of polyQ (Q31) peptides revealed a cross-β diffraction pattern typical of amyloid fibrils^24,25^. Other biophysical methods, including EPR, circular dichroism (CD), Fourier-transform infrared spectroscopy (FTIR), and UV resonance Raman (UVRR) spectroscopy of aggregated polyQ, indicated the formation of β-hairpins^26–32^. Recent studies of Htt filaments using electron cryo-tomography (cryo-ET) coupled with advances in missing-wedge compensation and subtomogram averaging (STA) have attempted to reveal more structural details. Branched and heterogeneous morphology of these filaments was revealed at low resolution, without molecular details^37,38^. Although the core of the Htt fibrils was not directly observed in these studies, they have helped to generate theoretical structural models of Htt filaments and guide the studies aimed at achieving this goal, including the work presented here. However, the absence of more detailed cryo-EM structures has precluded efforts to directly validate these models or guide more precise experiments to test current hypotheses on the structural basis underlying Htt aggregation, polymorphism, and toxicity.

Several fundamental questions about the structural properties of Htt filaments and the arrangement of the core of the fibrils remain unanswered. These include the following: 1) Are Httex1 fibrils twisted like other amyloids, or are they planar? 2) How are hairpins in individual protofilaments stacked, and how the straightness of fibrils is maintained? 3) How does polyQ length influence stacking and overall filament morphology? 4) What is the spatial distribution of the flanking domains, and how do they interact with each other and with the polyQ amyloid core? 5) How do the flanking domains influence the structure of the individual filaments and their lateral association and polymorphism?

Here, we present the first attempt to directly investigate the structural properties of in vitro-generated Httex1 fibrils by performing cryo-EM single-particle analysis (SPA). Our results and analyses showed that WT and mutant Httex1 form non-twisted filaments. 2D class averages and nonhelical 3D reconstructions provided new structural details and insights into the sequences of hairpins that form the core of the filaments. We found unpredicted stacking of β-hairpins and supramolecular polymorphs formed by the association of a larger number of protofilaments. We provide new insight into the effects of polyQ length and N-terminal domains on the structure of Httex1 fibrils and the arrangement of the polyQ flanking domains (first 17 N-terminal amino acids, Nt17, and C-terminal proline-rich domain, PRD) on the surfaces of the filaments. These findings broaden the current knowledge regarding the structure of fully assembled Httex1 filaments and provide valuable information that could guide future cryo-EM and structural studies on filaments derived from Htt and other disease-associated polyQ-containing proteins.

## Results

### Cryo-EM analysis of Httex1 filaments

To gain insight into the structure of filaments formed by Htt fragments and other polyQ peptides and proteins, we sought to determine the cryo-EM structure of exon 1 of the Htt protein (Httex1). Several preparations of Httex1 filaments were screened first by negative-stain transmission electron microscopy. These included filaments formed by Httex1 with polyQ lengths below (23Q) or above the pathogenic threshold of 36Q (43Q) **(Fig. 1)**. To break thick clusters of filaments and facilitate their uniform distribution in the vitreous ice layer, Httex1-43Q filaments were sonicated before application to the cryo-EM grid.

**Fig. 1.**
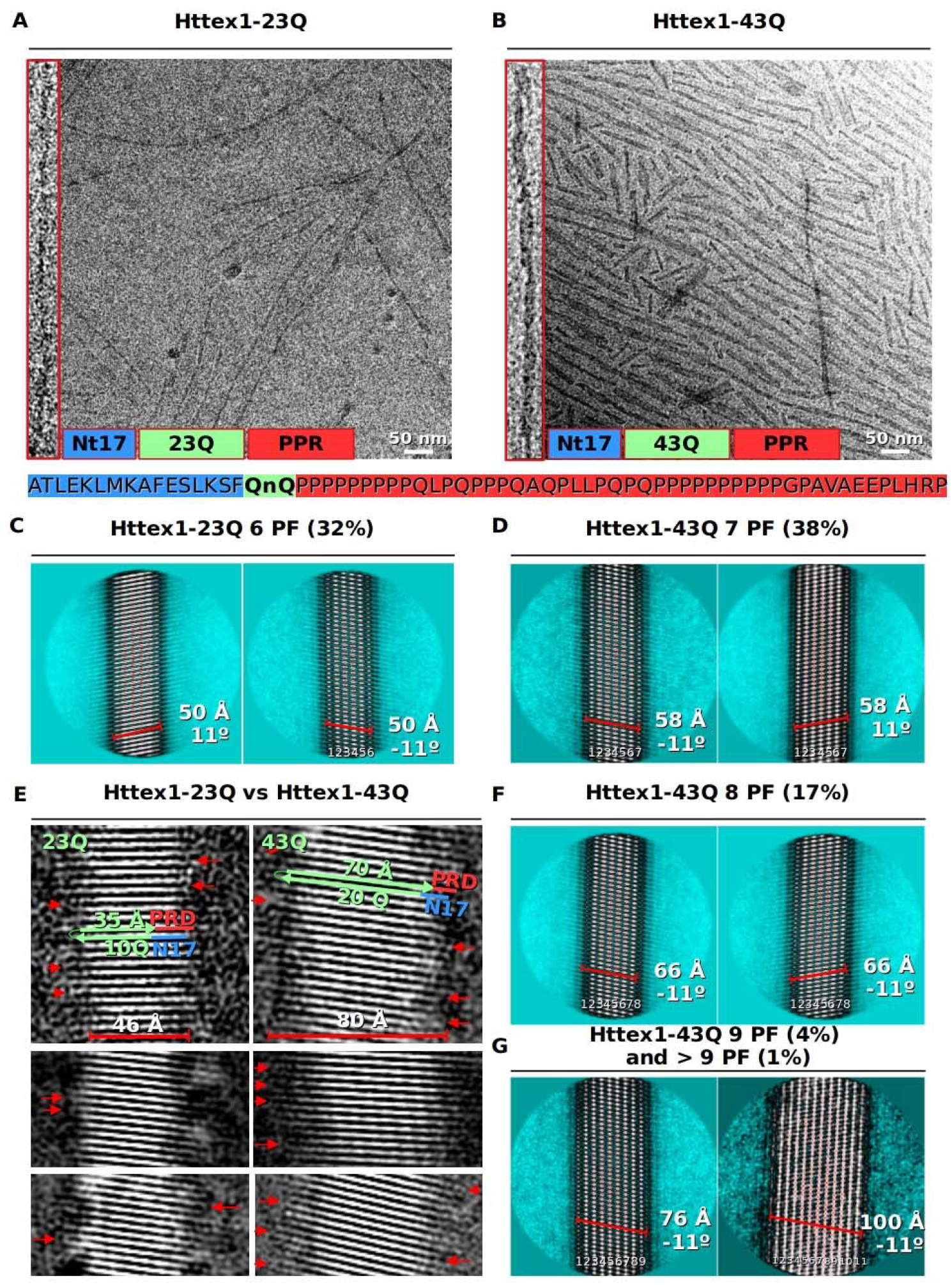
Cryo-EM micrographs of (A) Httex1-23Q and (B) Httex1-43Q filaments with an enlarged view of individual fibrils (left panel). Representative 2D class average images of major conformations of (C) Httex1-23Q (6 protofilaments) and of (D) Httex1-43Q (7 protofilaments). Representative 2D class average images of minor conformations of Httex1-43Q composed of (F) 8 protofilaments and of (G) 9 or more protofilaments. (E) 2D class average images of Httex1-23Q filaments (left panel) and Httex1-43Q (right panel) with variable β-strand stacking angles. Resolved β-turns characteristic of β-hairpins are highlighted with red arrows.

Inspection of a large quantity of raw cryo-EM images demonstrated that filaments of Httex1-23Q are tubular and, on average, longer than the planar ribbon-like filaments formed by the Httex-43Q protein **(Fig. 1A, B)**, in agreement with previous AFM studies^39^. The 17 residues closest to the N-terminal end (Nt17) and the proline-rich C-terminal domains (41–90 for 23Q, 61–110 for 43Q) are both nonvisible on raw cryo-EM images, most likely due to their high flexibility **(Fig. 1A, B)**.

Several rounds of reference-free 2D classification revealed stacked polyQ β-strands without signs of filament twisting. Most of the 2D class average images exhibit unexpected and previously unreported tilted β-sheet stacking, with up to ± 11° inclination relative to the filament axis **(Fig. 1C-G)**. This observation pointed to another level of heterogeneity even along a single Httex1 filament, thus complicating further single-particle analysis (SPA).

2D class average images enabled the estimation of filament dimensions along stacked polyQ β-strands and perpendicular to them. For better visualization of these strands, several 2D class average images were bandpass filtered in the range of 4–10 Å **(Fig. 1E)**. Httex1-23Q filaments along β-strands are (46 ± 2 Å) wide. The polyQ β-strand has a rise of ⍰3.5 Å per glutamine residue; thus, a single 46-Å-wide Httex1-23Q β-strand accommodates up to 10 glutamine residues and partially resolved flanking domains **(Fig. 1E, left panel)**. Two neighboring strands are separated by an inter-strand distance of 4.85 Å, linked by small β-turns visible on 2D class average images **(Fig. 1E, Fig. S1A, red arrows)** and most likely represent 23Q β-hairpin.

Similar estimations for Httex1-43Q filaments suggest that the single 80 Å wide β-strand accommodates up to 20 glutamine residues, in addition to a very short, resolved fragment of flanking domains **(Fig. 1E, right panel)**.

The nonhelical nature of Httex1 filaments became apparent after inspection of 2D class averages of segments extracted with large box sizes (>700 Å, >1,000 Å). This technique allowed us to measure the crossover distance and corresponding helical twist **(Fig. S2A)**. None of the inspected 2D class averages exhibit any sign of a crossover, typical of many amyloids studied by cryo-EM. Additional cryo-EM data collection and SPA were conducted with Httex1-43Q filaments incubated with nanoparticles (NPs), which were shown previously to decorate the surfaces of the filaments and unmask their polymorphism under cryo-EM conditions. The resulting contrast enhancement can be used to visualize crossovers of filaments derived from different amyloid-forming proteins with distinct sequence and structural properties under cryo-EM conditions^12^. Several rounds of reference-free 2D classification of segments extracted from cryo-EM images of (Httex1-43Q+NP) filaments confirmed the absence of crossover and demonstrated the nonhelicity of Httex1 filaments **(Fig. S2B)**. As a control, cryo-EM images of α-synuclein and Aβ-40 amyloid filaments incubated with NP were inspected, and crossovers were easily detected after 2D classification **(Fig. S2C, D)**.

Reference-free 2D classification also revealed filament views perpendicular to β hairpins with different numbers of protofilaments, each composed of a single β-sheet. Each Httex1-23Q filament was composed of 6 protofilaments **(Fig. 1C)**. Httex1-43Q filaments showed more variability, from 7 to 9, with a small fraction of filaments exhibiting 10 and 11 protofilaments **(Fig. 1D, F, G)**. This morphological polymorphism clearly indicates significant lateral associations of individual planar protofilaments, induced by the large, exposed hydrophobic surface area of the hairpin stacks.

The polyQ-flanking N- and C-terminal domains became apparent only on 2D class average images, with a higher signal-to-noise ratio than on raw images **(Fig. 2A, C)**. However, even after averaging, these densities had a low contrast and fuzzy appearance compared to the high density of polyQ domains **(Fig. 2A, C red arrows)**. Such flexible domains complicated 2D classification by inducing a certain level of misalignment of the polyQ amyloid core. As hypothesized, but not directly observed in previous structural studies of Httex1 filaments^40,41^, was the uniform distribution of fuzzy flanking domains on both sides of polyQ stacks **(Fig. 2A, C)**. This finding clearly indicates N-terminal-to-N-terminal or C-terminal-to-C-terminal stacking of two neighboring β-hairpins to maintain the characteristic antiparallel arrangement, which has lower free energy than a parallel arrangement. In addition, the measured distance between two adjacent flanking domains is 26 ± 2 Å, which corresponds to 3 antiparallel stacked hairpins **(Fig. 2C)**. A small proportion (6%) of selected Httex1-23Q filaments exhibited inter-filament twinning driven by association of the flanking domains^41^ **(Fig. 2B)**.

**Fig. 2.**
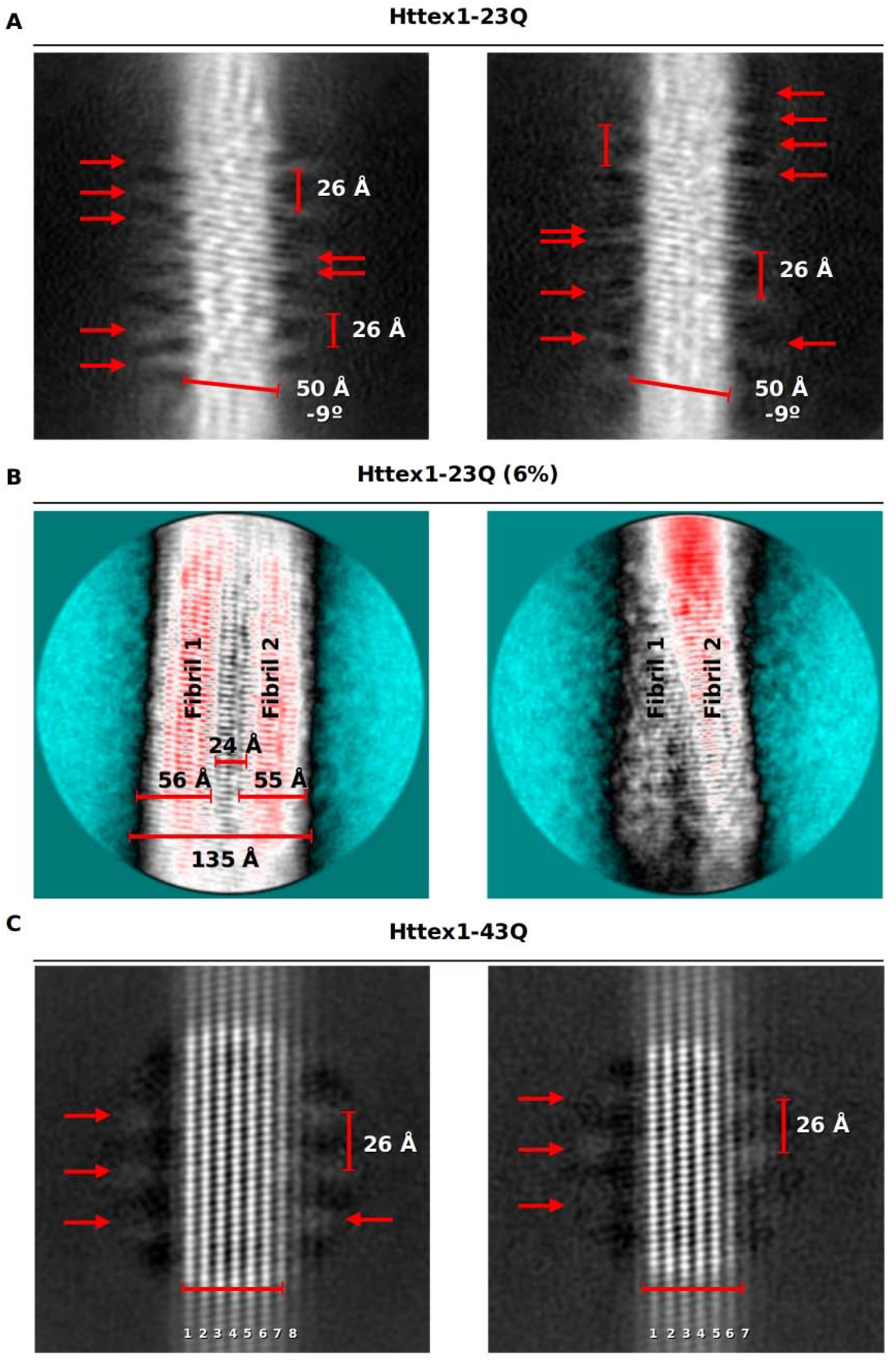
Representative 2D class average images of (A) Httex1-23Q and (C) Httex1-43Q filaments with resolved flanking domains (red arrows) distributed equilateral relative to the polyQ amyloid core. (B) A small proportion (6%) of selected Httex1-23Q filaments exhibits inter-filament twinning driven by association of the flanking domains.

The lack of helical symmetry significantly reduced the number of particles with different orientations needed for complete 3D reconstruction. Limited or nonuniform angular distribution of nonhelical filaments in vitreous ice usually leads to artifacts in 3D reconstruction, an inflated Fourier shell correlation (FSC) curve and consequent resolution estimation. Therefore, filament segments were extracted without overlapping and processed without helical symmetry application. Several useful features of helical processing were retained, such as limited angular search of out-of-plane (Tilt) and in-plane (Psi) rotation angles, for which the priors were estimated in a semiautomated selection procedure **(Fig. S3A, B)**.

Nonhelical 3D maps of Httex1-23Q and 43Q filaments were reconstructed **(Fig. 3)**, with estimated resolutions of 3.9 Å and 3.77 Å (FSC_0.143_ criterion) **(Fig. S4A, D)**. FSC curves showed significant oscillations in the range of 10–4 Å, indicating nonuniform angular distribution and overrepresentation of certain views. In addition, Httex1-43Q 3D reconstruction exhibits a certain degree of rotation angle misalignment, with a clear bimodal distribution **(Fig. S3E)**. Attempts to overcome misalignment did not improve the resolution and density appearance.

**Fig. 3.**
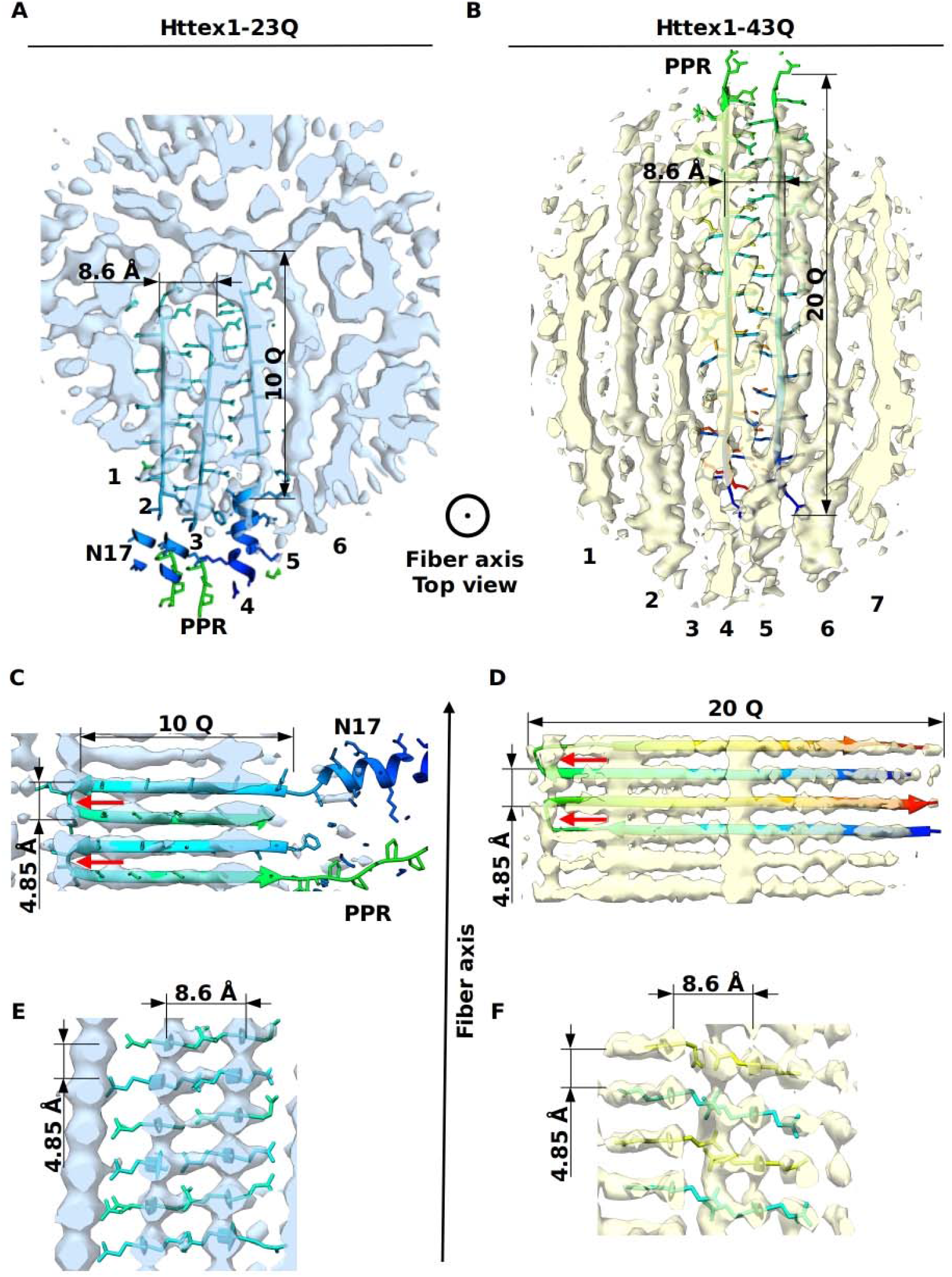
Central slices of the isosurface of asymmetrical 3D reconstructions with fitted homology-based models, top views of (A) the Httex1-23Q filament (FSC_0.143_= 3.9 Å) and of (B) the Httex1-43Q filament (FSC_0.143_= 3.77 Å). Orthogonal sliced side views of (C) and (E) Httex1-23Q filaments and of (D) and (F) Httex1-43Q filaments.

Notwithstanding these limitations, the resulting 3D reconstructions provide structural insights into the heterogeneous architecture of Httex1 filaments. The 3D reconstruction of Httex1-23Q filaments shows that they are formed by approximately six protofilaments separated by an inter-sheet distance of 8.6 Å **(Fig. 3A, C, E)**. The exact number of protofilaments is not clear due to the high flexibility of the peripheral protofilaments and the presence of fuzzy flanking domains. Each protofilament in the reconstruction is built from a stack of β-strands, with two adjacent strands separated by an inter-strand distance of 4.85 Å and linked by a β-turn of ∼3 glutamine residues **(Fig. 3C, red arrows)**, and represents a 23Q β-hairpin. The position of a given β-hairpin in a protofilament is produced by a vertical translation of 4.85 Å * 2 = 9.6 Å without rotation **(Fig. 3C)** or with a 180° vertical flip to account for antiparallel β-strand arrangement.

3D reconstruction of the most abundant (38%) 3D class of the Httex1-43Q filament indicates that it is built from 7 protofilaments separated by an inter-sheet distance of 8.6 Å **(Fig. 3B, D, F)**. Attempts to reconstruct Httex1-43Q filaments with larger numbers of protofilaments (8, 9, and 10) were not successful. Our data suggest that elongation of the polyQ domain (43Q vs. 23Q) straightens each protofilament and induces additional lateral associations.

Both Httex1-23Q and 43Q 3D reconstructions lack densities corresponding to horizontal β-arcs **(Fig. 3E, F; Fig. S1A, B (green arrows))**. This supports the model in which the turn of polyQ strands occurs within a β-sheet and not between two sheets of adjacent protofilaments.

The flanking terminal domains are not resolved in either of the 3D reconstructions due to their flexibility and ambiguity between the top and bottom views of filaments with tilted stacking. Protruding and flexible flanking domains may induce a strong preferential orientation of filaments in the ice layer and complicate further analysis. Preferential orientation of the filaments resulted in spurious artifacts in the 3D reconstruction, thus complicating correct interpretation and modeling of the reconstructed density.

Despite the high estimated resolution, 3D reconstructions did not allow complete interpretation and atomic model building de novo. Instead, a homology model was built **(Fig. 4A, B)** with the Httex1-44Q protein model from Boatz et al.^41^ as a template. Predicted Httex1-23Q and 43Q protein models were used for rigid-body fitting in 3D reconstructions, followed by real-space refinement with PHENIX **(Fig. 3)**.

**Fig. 4.**
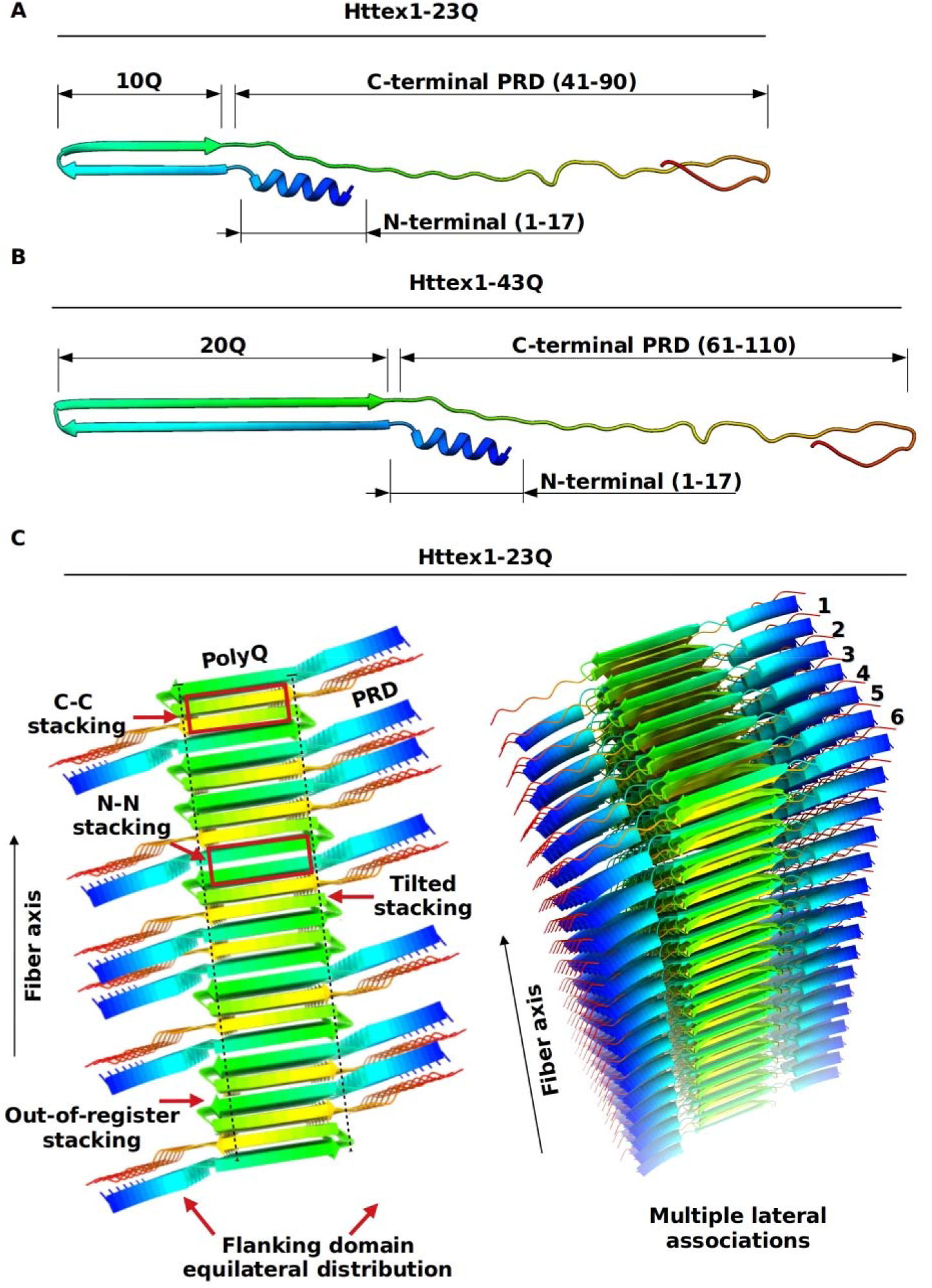
(A) Homology-based modeling of the protein structure of (A) Httex1-23Q and (B) Httex1-43Q filaments. (C) Schematic structural model shared by both Httex1-23Q and 43Q filaments, side view (left) and tilted view (right). Characteristic structural features are highlighted with red arrows, including β-strand tilted stacking with variable angles, out-of-register shifts, and equilateral distribution of flanking domains relative to the polyQ amyloid core.

Based on our findings, we propose a new Httex1 filament model, which complements previous models based on ssNMR studies^40,41^. The new model includes tilted stacking of β-hairpins and their misregistered states as well as the symmetrical distribution of the flanking domains relative to the polyQ core **(Fig. 4C)**.

Httex1 proteins with different polyQ repeat lengths (29Q and 37Q) were also screened with NS-EM and cryo-EM and rejected from further analysis because they form morphologies similar to those of Httex1-23Q and 43Q filaments **(Fig. S5A, B)**, with related complications in data processing.

### Effect of the Nt17 truncation on the structure of Httex1 filaments

Several studies suggest that the Nt17 domain strongly influences the structure and final morphology of the filaments formed by Httex1^9,42^. Together with the polyQ domain, they promote Httex1 aggregation in vitro and influence the kinetics of inclusion formation in cells. Previous studies from our group and others have also shown that Nt17 is exposed and flexible in Httex1 filaments, and recent studies from our group showed that its removal leads to a strong lateral association of the filaments with ribbon-like morphology^43,44^. Therefore, we further investigated the effect of the N-terminal truncation on the structure of Httex1 filaments.

We screened two mutant ΔNt17Httex1-23Q (Htt18-90-23Q) and ΔNt17Httex1-43Q (Htt18-90-43Q) filaments. Initial inspection of raw cryo-EM images followed by reference-free 2D classification of extracted segments revealed a high propensity of ΔNt17Httex1-43Q filaments to form lateral associations, which resulted in supramolecular polymorphism and formation of filaments with different numbers of protofilaments, similar to Httex1-43Q **(Fig. S5C)**. Therefore, the ΔNt17Httex1-43Q sample was excluded from further cryo-EM data collection and SPA.

Raw cryo-EM images demonstrated that ΔNt17Httex1-23Q filaments tend to bend. They resemble ribbons with variable width along a single filament **(Fig. 5A, red, yellow, and blue circles)**. Contrary to Httex1, the flanking C-terminal domains are visible already on raw images. Lateral associations of multiple protofilaments are less prominent for ΔNt17Httex1-23Q filaments, both on raw images and on 2D class averages, than Httex1-23Q filaments **(Fig. 5A, C)**.

**Fig. 5.**
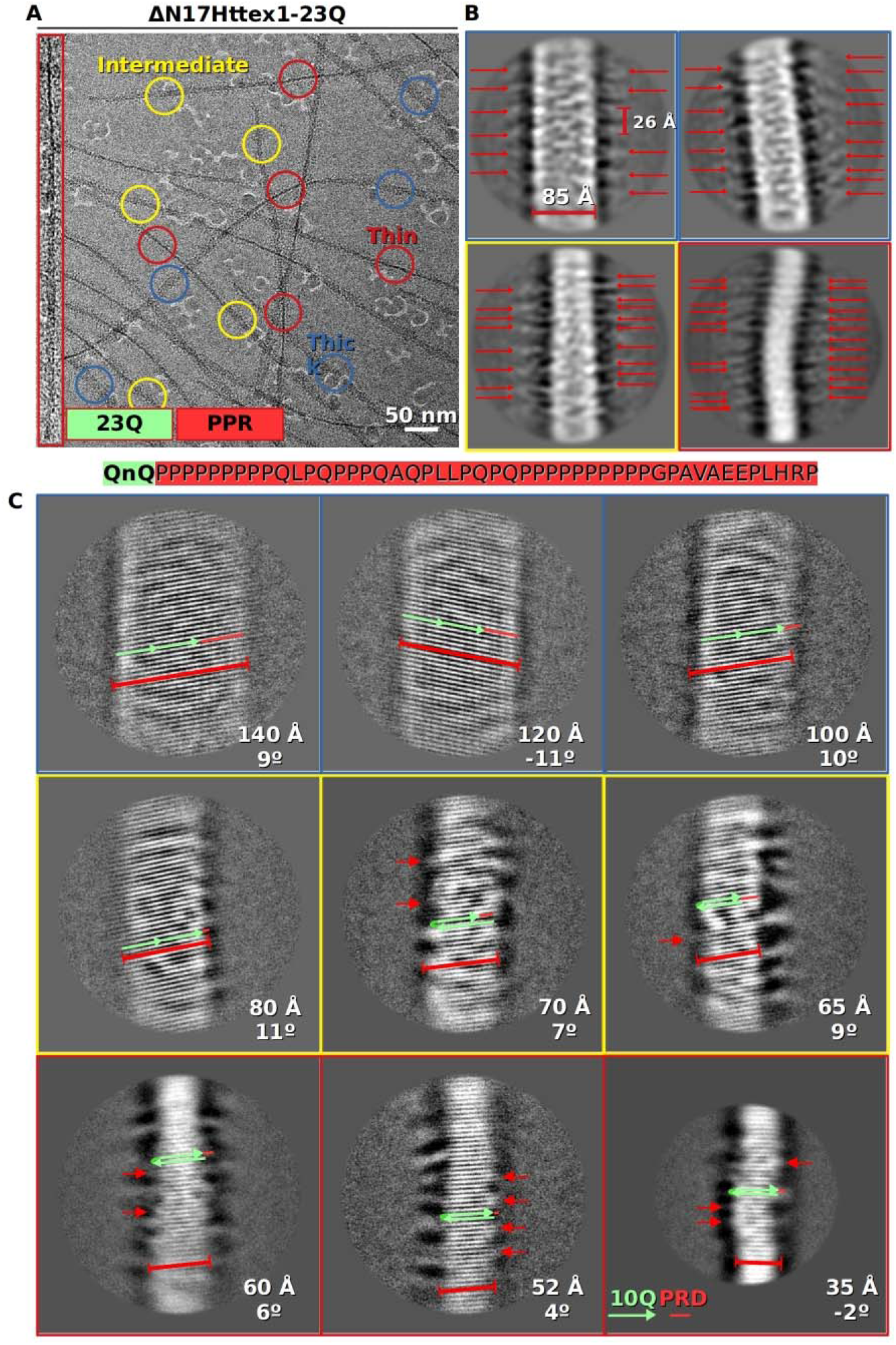
(A) Cryo-EM micrograph of ΔN17Htt-23Q filaments with an enlarged view of individual fibrils (left panel). (B) Representative 2D class average images of ΔN17Htt-23Q filaments, calculated with CTF amplitude correction only from the first peak of each CTF onward, for better visualization of flanking C-terminal domains (red arrows). (C) Representative 2D class average images of ΔN17Htt-23Q filaments with width and stacking angle variations. Resolved β-turns characteristic of β-hairpins are highlighted with red arrows.

Width variations along the same filament resembled crossovers and possible twisting **(Fig. 5A)**. Further 2D classification of particles extracted with large box sizes (>700 Å) revealed putative crossovers at 640 Å distance **(Fig. S1E)**. However, 3D refinement with helical symmetry application did not confirm the presence of twisting (not shown). For processing, we used a nonhelical cryo-EM SPA, which was similar to the pipeline used to process Httex1-23Q and 1-43Q, with the complications mentioned above.

Filament segments were re-extracted without overlapping and reprocessed without helical symmetry application. A limited angular search of out-of-plane and in-plane rotation angles around known priors was used. Single ΔNt17Httex1-23Q filaments were comprised of altering segments of different widths. Reference-free 2D classification revealed two major subpopulations of extracted segments, both with pseudo-helical β-strand stacking **(Fig. 5C).** Thin segments resemble horizontal stacks of β-strands, similar to full-length Httex1-23Q, but less straight and with a tendency to bend.

The thinnest segments are 35 ± 2 Å wide, in agreement with the size of the 23Q hairpin **(Fig. 5C)**. Similar to Httex1, small β-turns of ΔNt17Httex1-23Q hairpins were directly observed in 2D class average images **(Fig. 5C, Fig. S1C, red arrows)**. The thicker the segments, the more the β-strand stacks are prone to tilt, with up to a ±11° inclination perpendicular to the filament axis **(Fig. 5C)**. Strikingly, most strands in 2D class average images extend from 50 Å to 140 Å, far beyond the expected 35 Å width of one strand of 23Q hairpin **(Fig. 5C)**. To adopt such dimensions, polyQ β-strands should not form hairpins and would have to incorporate up to 20 amino acids from the C-terminal domain into elongated β-strands.

To visualize the flanking C-terminal domains, reference-free 2D classification with CTF amplitude correction only from the first peak of each CTF onward was used. The resulting 2D class average images revealed a distribution of flanking C-terminal domains on both sides of the polyQ domain, with a 26 ± 2 Å period and possible pairing of neighboring domains **(Fig. 5B, red arrows)**.

Despite the deletion of the exposed Nt17 domain, ΔNt17Httex1-23Q filaments also exhibit a non-uniform angular distribution in 3D. A limited number of orientations resulted in artifacts in 3D reconstructions of both thin and thick segments. Both 3D maps were interpreted only from the front plane along resolved β-strands. Sideway orientations perpendicular to hairpins were underrepresented; thus, the number of individual protofilaments that constituted a single filament was not resolved. 3D reconstruction of thin segments resolved along β-strands is a (60 ± 2) Å wide filament, suggesting that it is built from a stack of β-strands, 17 glutamine residues each with 5 unresolved glutamine residues and a protruding flanking C-terminal domain **(Fig. 6A)**. 3D reconstruction of 110 ± 2 Å thick segments from the same filament revealed different β-strand-based motifs. The most prominent is a four-stranded β-meander with 15 ± 2 Å strands of 5 glutamine residues **(Fig. 6B)**. Altogether, ΔNt17Httex1-23Q filaments represent an additional level of intra-filament structural heterogeneity compared to full-length Httex1-23Q filaments, which has not been previously reported for Htt proteins.

**Fig. 6.**
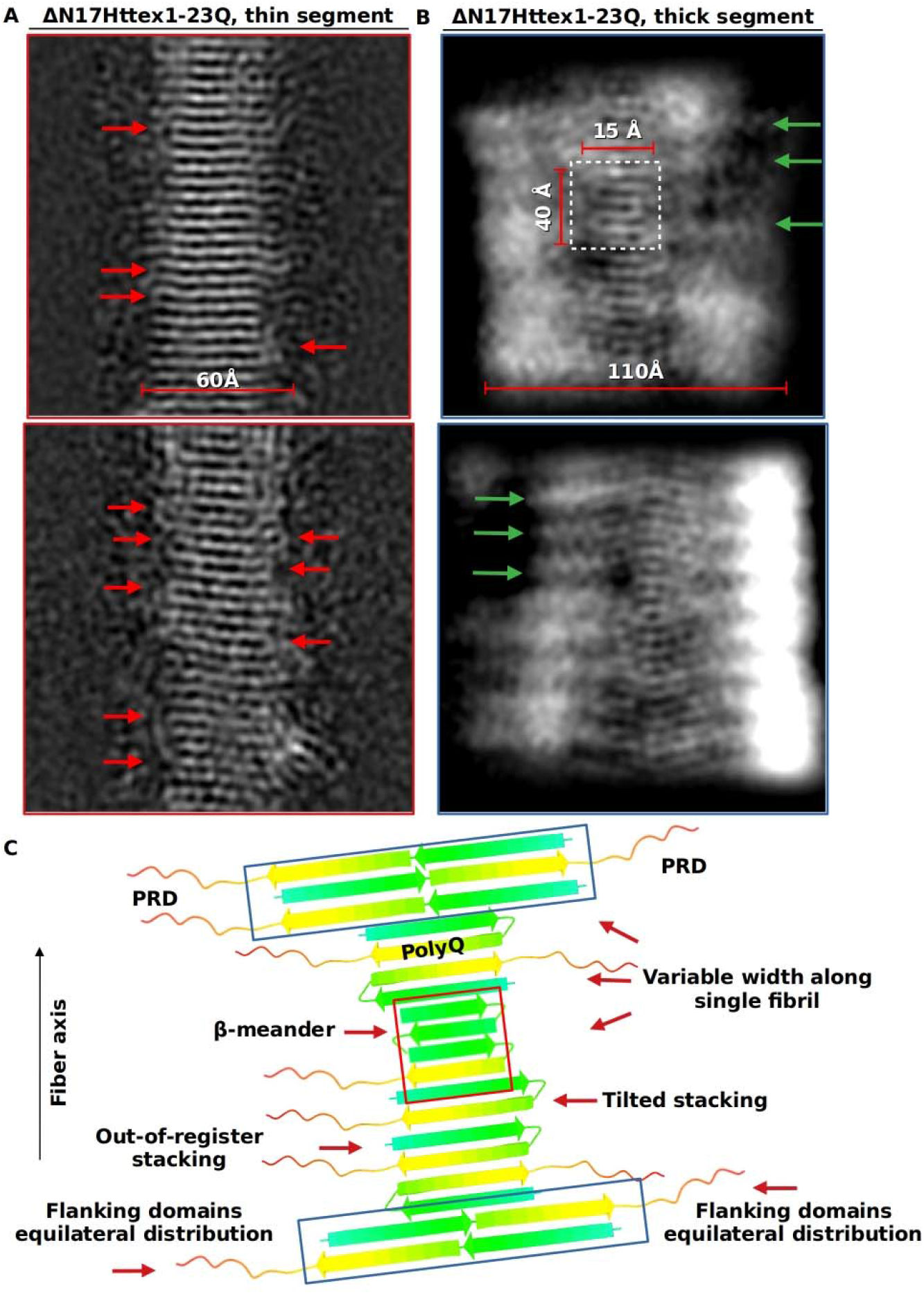
(A) Central slices of asymmetrical 3D reconstruction of (A) thin and (B) thick segments of the ΔN17Htt-23Q filament. Due to the strong preferential orientation, only the view along the β-strands is resolved. Resolved β-turns characteristic of β-hairpins of thin segments are highlighted with red arrows. The four-stranded β-meander is highlighted with a white square. Possibly resolved C-terminal proline-rich domains are highlighted with green arrows (top and bottom panels). (C) Schematic structural model outlining (red arrows) features characteristic of Nt17-truncated exon 1 filaments: variable single-filament width of single filament, mixture of β-hairpins, extended β-strands of variable width, and β-strand-based motifs including a four-stranded β-meander.

## Discussion

### 1 Complexity and heterogeneity of Httex1 filaments

Both Httex1-23Q and Httex1-431 filaments share a complex and heterogeneous architecture. These filaments are nonhelical, without observable helical crossovers characteristic of other amyloids such as the Aβ-40, αSyn, Tau and TDP-43 peptides^18–22^. These observations were confirmed by additional experiments with Httex1 filaments mixed with gold nanoparticles.

The short 23Q repeat, below the pathogenic 36Q threshold, resulted in the cylindrical appearance of Httex1-23Q filaments. Individual protofilaments of a single β-sheet are slightly bent, and bending increases for peripheral protofilaments (**Fig. 3A**). A more extended polyQ amyloid core (29Q, 37Q, and 43Q) of Httex1 filaments appeared to straighten the individual protofilaments and, together with multiple lateral associations, resulted in rectangular geometry (**Fig. 3B**), similar to that observed in recent studies of mutant Httex1-51Q filaments by cryo-ET subtomogram averaging^37^. Httex1-51Q filaments exhibit lamination and a large variation in width. In contrast, we did not observe narrow branching of Httex1-23Q and 43Q filaments.

Based on our cryo-EM analysis in 2D and 3D, we propose that the polyQ amyloid cores of both Httex1-23Q and 43Q filaments are built from stacks of β-hairpins. Each hairpin represents two antiparallel polyglutamine β-strands linked by small β-turns also composed of several glutamine residues. Cryo-EM allowed direct observation of such β-turns despite their small size and structural heterogeneity. In general, polyQ β-hairpins represent a particular case of hairpins that have not been previously observed in other amyloid filaments. In many structures of amyloid filaments recently solved by cryo-EM, β-sheets are organized in a well-aligned, in-register amyloid core^18–22^. A high energetic penalty prevents any misalignment of new β-strands on the growing filament. The polyQ amyloid core is formed by glutamine residues, and the β-hairpins are stabilized by glutamine side-chain hydrogen bonds, not via side chains of the polypeptide backbone, as in a conventional β-arc in many amyloids. Degenerate and thus promiscuous polyQ β-strands most likely account for the “hairy” bottle-brush appearance of Httex1 filaments, strongly marked in Httex1-23Q filaments. In addition, the tilted stacking of hairpins observed unexpectedly in this study had not been observed or suggested in previous models of Httex1 fibrils. Such tilting would be difficult if not impossible to detect with other structural methods. This finding may become a crucial point in understanding the structure of polyQ filaments. Tilted geometry raises the question of how neighboring polyQ β-strands are stacked and possibly indicates out-of-register states of “slippery” polyQ strands. The combination of tilting and out-of-register shifts provides a possible mechanism of filament straightness maintenance and avoidance of filament twisting by releasing the inherent steric stress of straight β-sheets.

Importantly, β-hairpins and proposed vertical stacking as the predominant architecture of both Httex1-23Q and 43Q filaments could be favored by slightly different protein constructs and different aggregation conditions. This is particularly important for the Httex1-23Q filaments for which previous structural studies predicted the β-arc and not the β-turn type of packaging.

The flanking N-terminal (Nt17) and C-terminal (proline-rich domain, PRD) domains of both filaments are only partially visible until after 2D classification, where the signal-to-noise ratio of 2D class average images increased relative to raw extracted particles. However, densities corresponding to flanking domains are weaker and less resolved than the polyQ amyloid core. These densities exhibit a characteristic fuzzy appearance, more prominent in the case of the Httex1-23Q filaments (**Fig. 2A**). Taken together, these observations indicate that flanking domains are flexible, with noticeable motion proximal to the polyQ core. The distribution of flanking domains around the filament axis appeared on average equilateral for both 23Q and 43Q filaments (**Fig. 2A, C**). This distribution is consistent with the antiparallel geometry of β-hairpins and indicates N-to-N or C-to-C stacking of adjacent hairpins within a single protofilament. In addition, the flanking domains appear to play an important role in the stabilization of the “slippery” polyQ core by preventing large out-of-register shifts^26^.

### 2 PolyQ-dependent lateral associations

Both the Httex1-23Q and Httex1-43Q protofilaments tend to undergo lateral associations, which appear to be driven by hairpin stacks of individual protofilaments. Large, exposed hydrophobic surface areas and hydrogen-bonding edges of β-hairpins induce the formation of compact bundles. The number of individual protofilaments in each bundle varies and depends largely on the size of the polyQ domain.

The predominant (38%) conformation of Httex1-23Q filaments is formed by six protofilaments. Certain flexibility of individual Httex1-23Q protofilaments resulted in a compact cylindrical shape, with outer density corresponding to bent peripheral protofilaments and fuzzy flanking domains. Such bending, if strong enough, should prevent the formation of an interface with a new straight β-sheet, potentially limiting the number of protofilaments in a single filament of Httex1-23Q. This observation explains why we did not observe Httex1-23Q filaments with more than 6 protofilaments.

Elongation of the polyQ domain noticeably straightens the Httex1-43Q protofilaments, increases hydrophobic surface area, and thus promotes stronger lateral associations. The major filament population is formed from seven (40%) protofilaments. Assemblies with eight (18%), nine (4%), ten (4%) and eleven (1%) protofilaments are also present. Bundles with a smaller number of protofilaments are most likely less stable and tend to form associations with the exposed hydrophobic surface of nearest protofilaments. Assemblies with large numbers of protofilaments (>11) may exist but may have been divided into smaller fragments during sonication or became too thick to fit into the vitreous ice layer.

As predicted in previous studies^26,40,41^, the flanking domains in both Httex1-23Q and Httex1-43Q filaments likely play an important role in determining the number of protofilaments within the fibril. Equilateral distribution of these domains and degrees of motion relative to the amyloid core can potentially prevent association with a new protofilament, either by electrostatic repulsion or by mechanical blockage of the exposed surface of the protofilament. Our results also suggest another type of inter-filament association based on possible intertwining of flanking domains **(Fig. 2B)**. However, only a small 6% fraction of 2D class averages of Httex1-43Q filaments demonstrate such intertwining. The distance between two intertwined filaments is 25 ± 2 Å, which is too small to accommodate the flanking domains of two filaments.

### 3 Intra- and inter-filament heterogeneity

The unique architecture of Httex1 filaments may also reflect a high degree of structural heterogeneity, even within the single filament.

First, tilted stacking of polyQ β-hairpins with variable stacking angles represents a novel structural feature of polyQ filaments. The hairpin stacking angle varies continuously from a horizontal 0° to ± 11° inclined arrangement. Stacking angle variations are observed even within the single protofilament and are based on variable out-of-register shifts of β-hairpins in both directions toward the N-terminus or C-terminus. Larger stacking angles and thus larger out-of-register shifts are most likely prohibited by the presence of flanking domains. Prolines from the C-terminal domain cannot adopt the extended β-sheet conformation. The large positively charged side chain of lysine from the N-terminal domain (ATLEKLMKAFESLKSF) also limits out-of-register shifts due to the large free-energy penalty of charge desolvation^26,45^.

Second, Httex1-43Q filaments exhibit heterogeneity in the number of protofilaments constituting a single filament. 2D class average images of segments extracted from the same Httex1-43Q filament exhibit different numbers of protofilaments. This observation clearly indicates that Httex1 filaments, especially those with long polyQ domains, can grow in two dimensions: along the filament axis by antiparallel stacking of hairpins and perpendicular to the filament axis by lateral associations. Most likely, this heterogeneity is characteristic of Httex1 filaments with large polyQ domains (>36Q). Third, there was intra-filament heterogeneity from the distribution of flanking domains relative to the filament axis. On a large scale, flanking domains are distributed equilateral to the filament axis. No preferential orientations of flanking domains were observed. However, at the level of individual hairpins, the orientation of neighboring hairpins is random. β-Strands of adjacent hairpins are stabilized by glutamine side-chain hydrogen bonds, either N-to-N or C-to-C, with equal probability **(Fig. 4C)**.

Truncation of the flanking Nt17 domain resulted in additional types of heterogeneity. The ΔNt17Httex1-23Q filaments exhibit striking width variations, from 35 Å to 120 Å. Thin regions appeared flexible, suggesting that the polyQ amyloid core assembled differently from straight Httex1-23Q filaments. The width of most segments exceeds the expected 35 Å width of the 23Q hairpin. One possible explanation is that N-terminal domain truncation lowers the free energy and geometric restrictions of full-length Httex1 filaments and breaks the characteristic β-hairpin architecture in most regions within a single filament. In order to adopt variable dimensions, β-strands must somehow incorporate up to 20 amino acids from the C-terminal domain to achieve sufficient length. However, proline residues are unable to adopt β-sheet structures due to space and hydrogen-bonding limitations. Potentially, thick segments could represent two β-sheets zipped together; however, we did not observe any visible interfaces between them.

The longer the β-strands, the higher the tilting angle is, with a maximum at ± 11°. The maximum tilt angle of 11° is the same for the Httex1-23Q and ΔNt17Httex1-23Q filaments, which indicates a role of the C-terminal domain in preventing larger tilts. Variable tilting angle is likely to be a universal compensation mechanism for both full-length and N-terminally truncated Httex1 filaments. Surprisingly, Nt17 truncation significantly removes inter-fiber heterogeneity, since ΔNt17Httex1-23Q filaments lack detectable lateral associations characteristic of Httex1 filaments; however, the mechanism behind Nt17 involvement in lateral associations remains obscure. From our observations, a single ΔNt17Httex1-23Q filament is composed of a small number of protofilaments, or even one planar β-sheet, and is heterogeneous along its length. Since we were not able to resolve 3D reconstructions of ΔNt17Httex1-23Q in the direction perpendicular to β-strands, it is possible that Nt17 truncation locally induces different types of packaging along a single filament. Local packaging changes may appear as β-strand width variations.

Altogether, these observed types of heterogeneity seem to be characteristic of polyQ filaments and impose strict limitations on the usage of the cryo-EM helical reconstruction method.

It should be noted that differences in some of our observations in relation to previous studies could be because the polymorphism of Httex1 fibrils is highly influenced by 1) solution conditions such as protein concentration, pH, buffer components, etc.; 2) small variations in the protein sequence; or both. Our findings here were based on filaments derived from untagged Htt proteins (native sequence) aggregated under specific conditions, whereas many of the previous studies were based on polyQ-peptide models or Httex1 proteins containing additional nonnative sequences (e.g., histidine tags or S tag KESAAAKFERQHMDS) ^15,26,40,41,46^. Furthermore, changes in solution incubation conditions may force different Httex1 filament structural conformations and types of polymorphisms. For example, β-hairpin formation could be a function of polyQ length and aggregation conditions, which in some cases favor β-arc architecture or a mixture of both filaments or within a single filament. This underscores the importance of continuing to explore different conditions to generate fibrils with distinct morphologies and solve their structures with cryo-EM.

### 4 Conclusions

Cryo-EM is a powerful structural technique capable of solving the structures of amyloids in near-native conditions when coupled with modern algorithms. Our ability to resolve individual β-strands along the filament axis combined with helical symmetry peculiar to many amyloid filaments resulted in atomic models with large predictive potential.

Polyglutamine huntingtin filaments represent a unique type of amyloid. They lack helical symmetry, showing strong intrinsic heterogeneity between individual filaments and, more importantly, within the single protofilament. Our cryo-EM studies provide new insights into the structural organization of the Httex1 monomers within preassembled filaments, reveal several unique structural features, and refine our understanding of how the filaments form, albeit not at the atomic level.

Despite not achieving atomic resolution, the insight our 3D structures provide expands our knowledge of Httex1 and polyQ-driven amyloid formation and will help guide future studies to investigate and target the mechanisms of Htt filament formation and disassembly. Direct observation of the polyQ stacking and flanking domain distribution of “short” Httex1-23Q and “long” Httex1-43Q filaments revealed their potential weak points. Based on our updated model, loops connecting two β-strands of the same hairpin are exposed to the solution and accessible for potential drug molecules. Disruption of these loops could remove constraints of β-hairpins and could lead to either complete Htt filament disassembly or a noticeable shortage. It should be noted that due to inherent intra-filament heterogeneity, these loops are exposed differently to the solution and could also be partially hidden by flanking domains.

The high propensity of Httex1 protofilaments to undergo lateral associations and our hypothetical structural model underpinning this process suggest another weak point in Htt filament assembly. Potential drug molecules should target the hydrophobic interface formed by β-sheets of neighboring protofilaments to prevent attachment of new protofilaments.

We expect that many members of the family of polyglutamine filaments share a similar structure and intrinsic heterogeneity. Due to the lack of helical symmetry, future efforts should focus on shortening long preassembled filaments, either biochemically or by mechanical methods, while preserving the underlying hairpin-based architecture. Optimized samples should be processed with a conventional single-particle approach with short huntingtin filaments as “single particles.” Intrinsic intra-filament heterogeneity can be bypassed by utilizing 3D focused classifications, as well as by local symmetry 3D refinement. Potential improvements in structure determination may come from tilting data collection schemes or from hybrid approaches combining single-particle analysis and electron cryo-tomography.

## Materials and methods

### Filament preparation

Huntingtin exon 1 filaments in full-length and N-truncated form (without the first 17 amino acids) were prepared according to a previously described protocol^47^. Briefly, Httex1 with 23Q, and 43Q were prepared using fused to a SUMO tag^47^ and Httex1 29Q, 37Q as well as the two mutant Htt18-90 with 23Q and 43Q were fused to an Intein^48^ tag. Both methods enable the production of native tag-free huntingtin exon-1 monomer. Then, for the filament’s preparation monomers were disaggregated as previously reported^47^ and resuspended in PBS buffer and the pH was adjusted to 7.4. Finally, the filaments were formed by incubating the solution of monomers at 37°C without shaking.

### Cryo-EM grid preparation and data collection

Recombinant filament samples were screened with negative staining (NS) EM for filament concentration and morphology. Aliquots of 3 µL of optimized filament samples were applied onto glow-discharged, 300-mesh copper Quantifoil R2/1 or R1.2/1.3 grids. Cryo-EM grids were blotted from both sides with a Vitrobot Mark IV (Thermo Fisher Scientific: TFS) operated at 95% humidity and 20 °C and plunge frozen in liquid ethane cooled by liquid nitrogen. Cryo-EM grids were screened for high purity, concentration and homogeneity of the filaments and the presence of a thin layer of vitreous ice and stored in liquid nitrogen for further imaging. Homogeneous distribution of the filaments within a hole of the carbon grid allowed faster and more effective data acquisition with beam tilt implementation.

A representative number of cryo-EM images were collected on an FEI Tecnai F20 200 kV transmission electron microscope (TEM) equipped with a FalconIII direct electron detector (TFS) at CIME, EPFL, and subjected to filament tracing followed by reference-free 2D classification on the GPU cluster at EPFL. The resulting 2D class averages with original cryo-EM images were included in a beam time application at the Center for Cellular Imaging and Nanoanalytics (C-CINA), Basel, Switzerland.

For determination of fibril structure, samples were imaged on a TFS Titan Krios TEM operated at 300 kV and equipped with a Gatan Quantum-LS imaging energy filter (GIF, 20 eV energy loss window; Gatan Inc.), or on a TFS Glacios Cryo-TEM (200kV). On the Titan Krios TEM were acquired on K2 Summit electron counting direct detection camera (Gatan Inc.) in dose fractionation mode (40 frames) using Serial EM software^49^ at a magnification of 100,000× (physical pixel size 1.058 Å) and a total dose of 48 electrons per square angstrom (e^-^/Å^2^) for each exposure. On the Glacios, images were acquired on a K3 electron counting direct detection camera (Gatan Inc.) in dose fractionation mode (40 frames) using Serial EM software at a magnification of 100,000× (physical pixel size 1.05 Å) and a total dose of 49 electrons per square angstrom (e^-^/Å^2^) for each exposure. Micrographs were drift corrected and dose weighted using MotionCor2^50^. The contrast transfer function (CTF) for each micrograph was estimated using CTFFIND4.1^51^ [44] through the focus interface [45]. After manual inspection through the focus interface, the best 4250 (Httex1-23Q and 43Q) filaments and the best 2744 (ΔNt17Httex1-23Q) cryo-EM images were transferred to local storage for further processing on a GPU cluster (4 x NVIDIA Tesla V100).

### Image processing

Pseudo-helical reconstruction was carried out with RELION3.1^52^ or with cisTEM^53^. Several hundred representative nonoverlapping filaments were manually selected using the e2helixboxer.py from EMAN2^54^. Dose-weighted averages were denoised with a pretrained JANNI network^55^ and subjected to semiautomated filament tracing with the crYOLO neural network^56^. Filament start and end coordinates in STAR format were imported into RELION3.1, and 1,052,700 Httex1-23Q segments, 1,955,000 Httex1-43Q segments and 592,815 ΔNt17Httex1-23Q segments were extracted without overlapping using a box size of 256 pixels. Extracted segments were subjected to several rounds of reference-free 2D classification to clean up the initial dataset and select only particles with clear 4.8 Å β-strand separation along the filament axis.

Further processing was performed without helical symmetry application. For convenience, several features of helical processing were maintained, such as limited angular search of out-of-plane (AngleTilt) and in-plane (AnglePsi) rotation angles, whose priors were calculated during filament tracing by crYOLO **(Fig. S3)**.

Cryo-EM images of amyloids incubated with nanoparticles (NPs) were collected by Cendrowska et al. and processed as described previously.

2D class averages of Httex1-23Q filaments were inspected, and suspicious classes with fuzzy periphery were removed from further processing. 2D class averages of Httex1-43Q filaments revealed polymorphs with different numbers of protofilaments (7–11). Such 2D class averages were identified and processed separately. Since different polymorphs of Httex1-43Q filaments with variable numbers of protofilaments share the same appearance in 2D class averages inspected from the front, several rounds of 3D classifications with a tight mask including only the central region of the filaments were needed to further separate polymorphs.

Since the flanking domains of Httex1-43Q were not resolved after 2D classification in RELION3.1, cisTEM was used for 2D classification of these filaments.

Initial models were generated de novo with RELION3.1 for Httex1-43Q filaments or with cisTEM for Httex1-23Q filaments. Rounds of asymmetrical focused 3D classifications with a mask including the central region of the Httex1-43Q filament were iterated with 3D auto-refinement, resulting in a 3D reconstruction with an estimated global resolution (FSC_0.143_ criterion) of 3.77 Å **(Fig. S4D)**. However, FSC curves show significant oscillations in the range of 10–4 Å, which is an indication of a non-uniform angular distribution and overrepresentation of certain views. In addition, the rotation angle demonstrated a bimodal distribution **(Fig. S4E)** resulted in certain degree of misalignment. Attempts to use only particles from the major peak did not improve the overall resolution or appearance of the 3D reconstruction.

Particles constituting the best 2D class averages of Httex1-23Q filaments were combined into a single stack and imported to cisTEM. Additional rounds of 2D classifications followed by 3D auto-refinement and 3D manual refinement (3D classification) with a tight mask around the central region of the Httex1-23Q filament resulted in a 3D reconstruction with an estimated global resolution (FSC_0.143_ criterion) of 3.9 Å **(Fig. S4A)**.

Despite a high estimated resolution, 3D reconstructions did not allow complete interpretation and atomic model building de novo. Instead, homology-based modeling with SWISSMODEL was used, with a theoretical model built for the Httex1-44Q protein with EM and NMR constraints as a template^41^. The resulting Httex1-23Q and 43Q protein models were used for rigid-body fitting into cryo-EM 3D reconstructions **(Fig. 3)** followed by several rounds of real-space refinement with PHENIX^56^.

## Data availability

All 3D reconstructions will be provided by request. Raw cryo-EM videos will be uploaded to the Electron Microscopy Public Image Archive (EMPIAR).

## Acknowledgments

This work was supported by funding from CHDI. We thank Mohamed Chami, and Lubomir Kovacik (BioEM facility, Biozentrum, Basel, Switzerland) with data collection on the TFS Titan Krios and TFS Glacios. We are grateful to Prof. Patrick van der Wel (Zernike Institute for Advanced Materials, University of Groningen, Netherlands) and Ricardo Guerrero-Ferreira (Robert P. Apkarian Integrated Electron Microscopy Core, Emory University, USA) for critical reading of the manuscript and their constructive and valuable feedback and input.

