## Supplementary data for "The structural basis of huntingtin (Htt) fibril polymorphism, revealed by cryo-EM of exon 1 Htt fibrils"

Key words: Huntington's disease, Huntingtin, Fibrils, Cryo-EM, Polymorphism

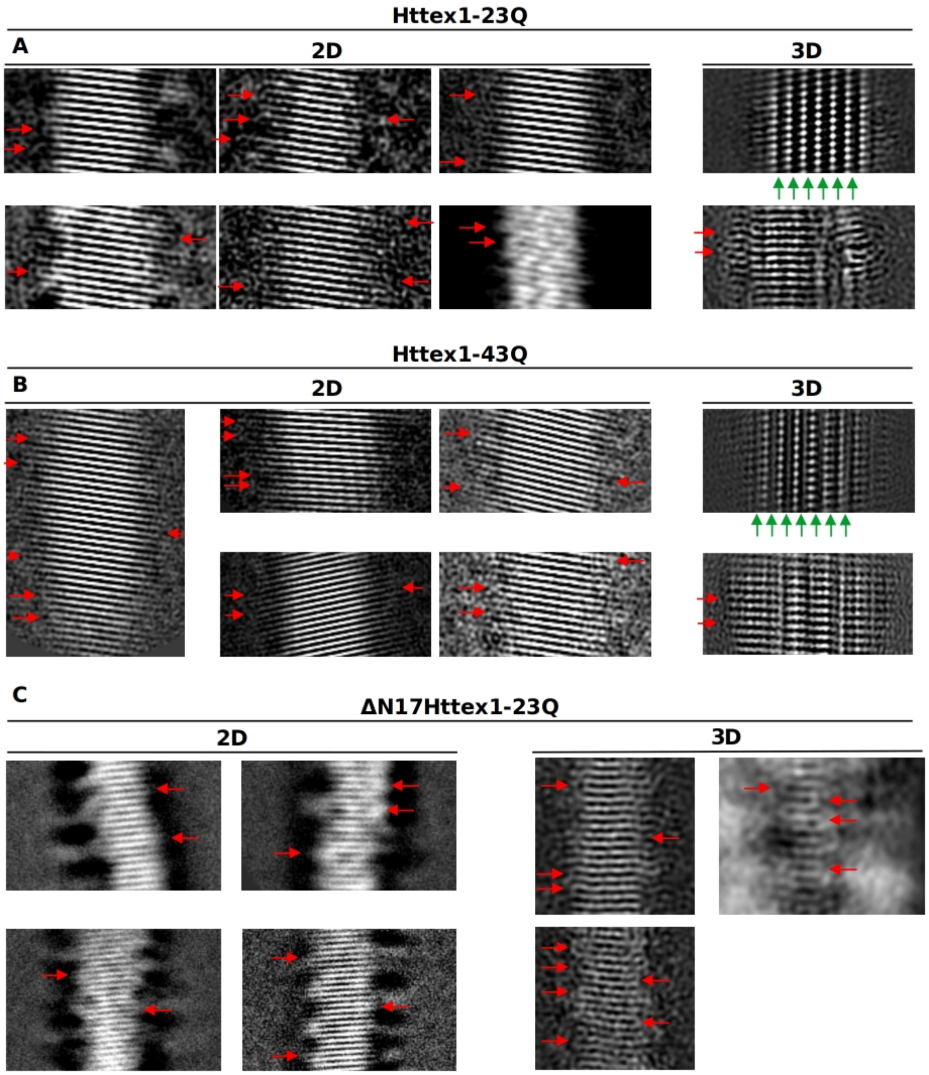


**Fig. S1.** (A) Representative 2D class average images of Httex1-23Q filaments (left). Resolved β-turns characteristic of β-hairpins are highlighted with red arrows. Orthogonal slices along and perpendicular to β-strands of 3D reconstruction of Httex1-23Q filament (right). The lack of β-arcs is highlighted with green arrows. (B) Representative 2D class average images of Httex1-43Q filaments (left). Resolved β-turns characteristic of β-hairpins are highlighted with red arrows. Orthogonal slices along and perpendicular to β-strands of 3D reconstruction of the Httex1-43Q filament (right). The lack of β-arcs is highlighted with green arrows. (C) Representative 2D class average images of ΔNt17-Httex1-23Q filaments (left). Resolved β-turns characteristic of β-hairpins are highlighted with red arrows. Slices along β-strands of 3D reconstruction of ΔNt17-Httex1-23Q filament (right).

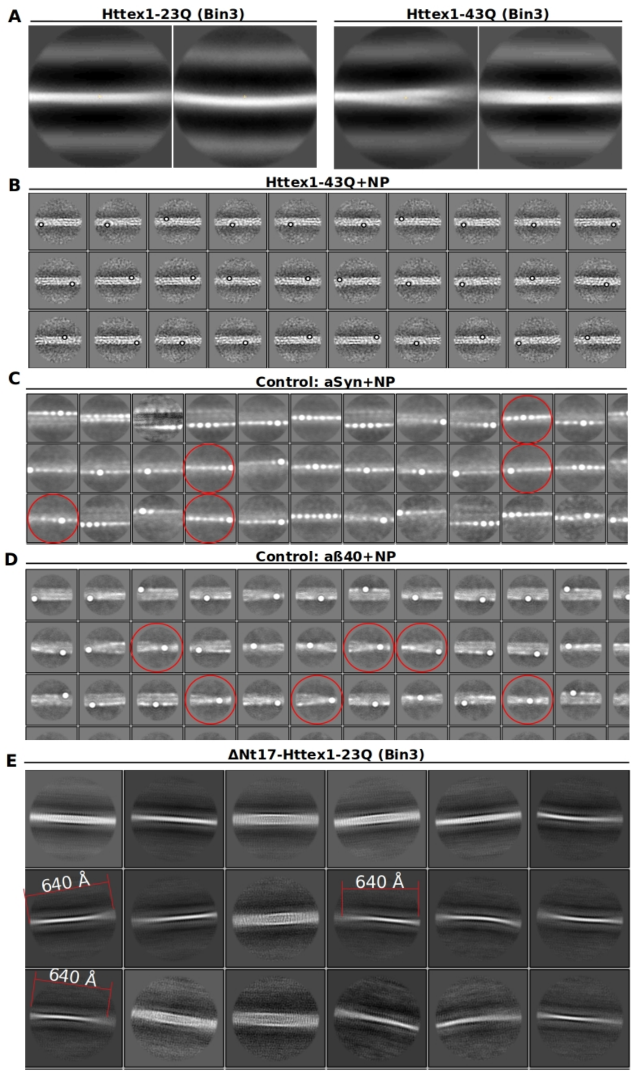


**Fig. S2.** (A) Representative 2D class averages of large extracted segments (700 pixels) from cryo-EM micrographs of (left) Httex1-23Q and (right) Httex1-43Q filaments, without detectable crossover. (B) Representative 2D class average images of Httex1-43Q incubated with gold nanoparticles confirm the lack of twisting of Httex1 filaments. Representative 2D class average images of segments extracted from previously collected cryo-EM micrographs (by Cendrowska et al.)[12] of (C) wild-type α-synuclein and (D) Aβ-40 filaments incubated with gold nanoparticles. Resolved crossovers are highlighted with red circles. (E) Representative 2D class average images of large (700 pixels) extracted segments from cryo-EM micrographs of ΔNt17-Httex1-23Q filaments. An unconfirmed 640 Å crossover is highlighted.

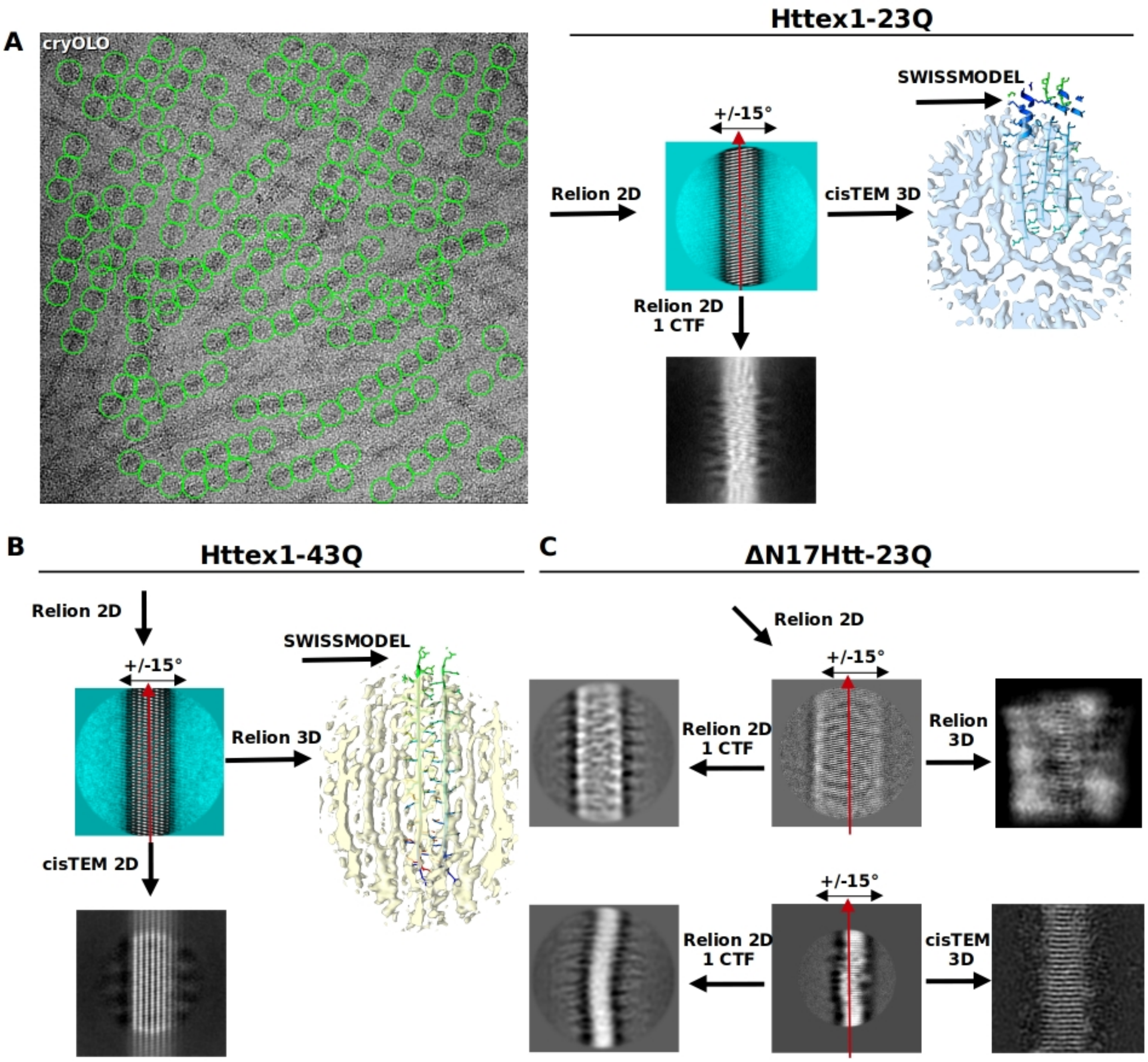


**Fig. S3.** Cryo-EM single-particle analysis scheme. (A) Representative cryo-EM micrograph of Httex1-23Q filaments with non-overlapped segments picked by crYOLO (left) and scheme of 2D classifications 3D classifications and refinement (right). Scheme of 2D and 3D processing of (B) Httex1-43Q and (C) ΔNt17-Httex1-23Q filaments.

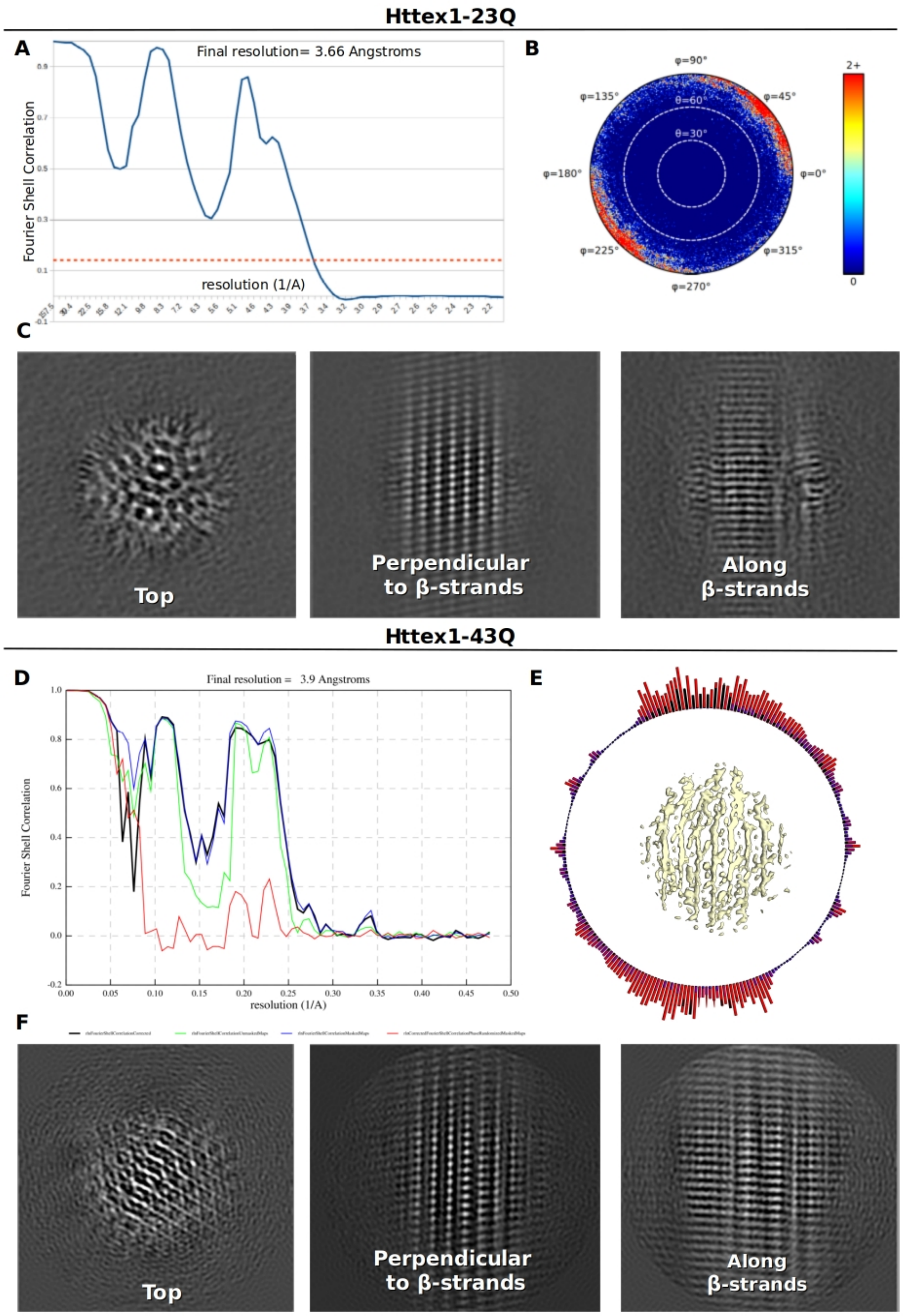


**Fig. S4.** Fourier shell correlation (FSC) curves of 3D reconstructions of (A) Httex1-23Q and (D) Httex1-43Q filaments. In-plane rotation angle distribution of the (B) Httex1-23Q filament from cisTEM and of the (E) Httex1-43Q filament from RELION. Orthogonal slices (top, perpendicular to β-strands and along β-strands) through 3D reconstructions of (C) Httex1-23Q and (F) Httex1-43Q.

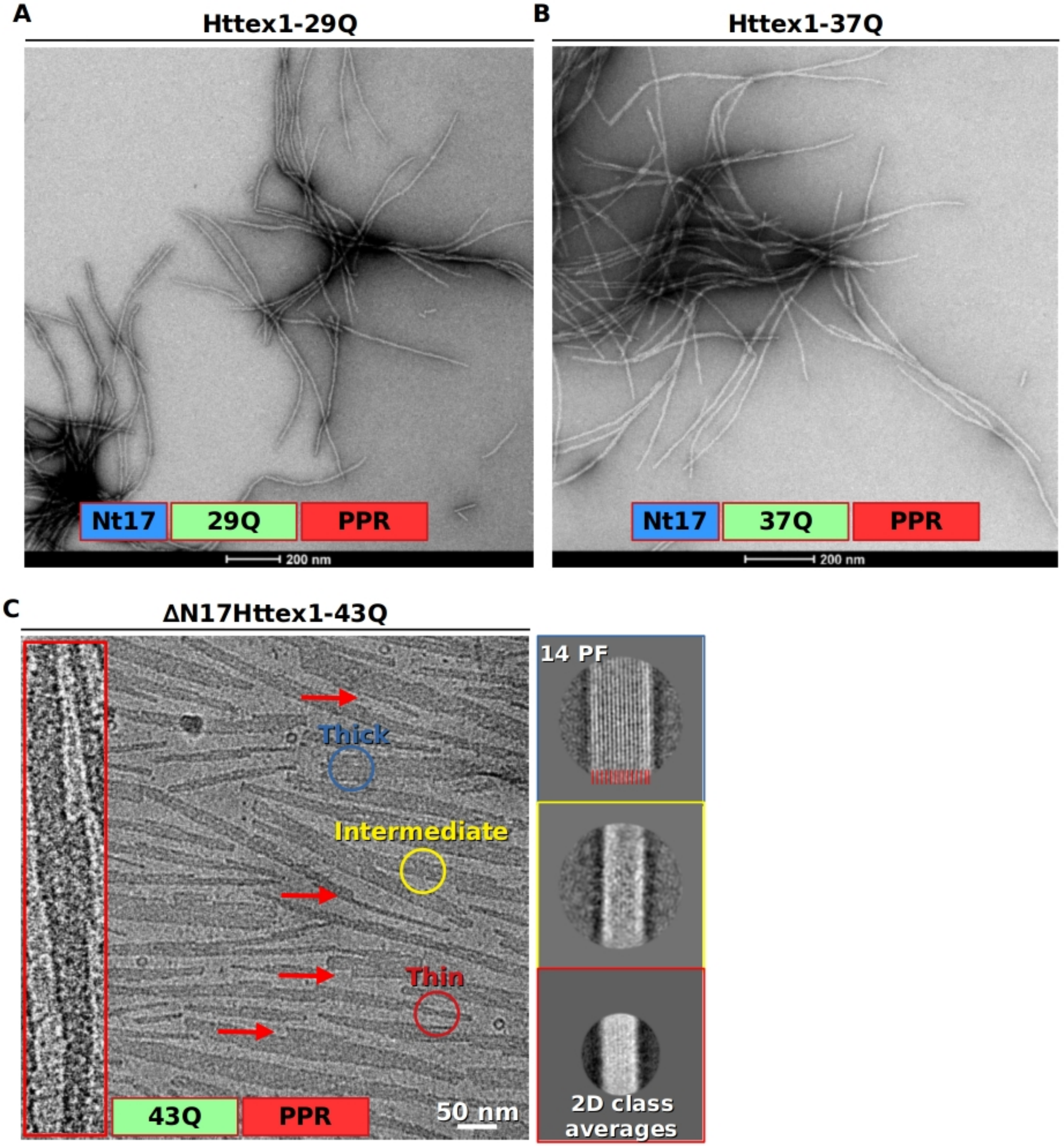


**Fig. S5.** Micrographs of Htt filaments excluded from single-particle analysis, with different domain compositions and polyQ lengths. Negative staining micrograph of (A) Httex1-29Q and (B) Httex1-37Q filaments. (C) Cryo-EM micrograph of N17-truncated ΔN17Httex1-43Q filaments with an enlarged view of an individual filament with multiple lateral associations (left panel). Representative 2D class average images of ΔN17Httex1-43Q filaments (right).
